# Application of the ORBIT Protocol for Targeted Gene Deletion to *Mycobacterium abscessus*

**DOI:** 10.64898/2026.10.23.761380

**Authors:** Nikita H. Amin, Timothy W. Sherrier, Nancy A. Woychik

## Abstract

*Mycobacterium abscessus* (Mab) is an emerging opportunistic pathogen characterized by extensive intrinsic drug resistance, underscoring the need for precise genetic tools to define mechanisms of pathogenicity and persistence. The ORBIT (<u>o</u>ligonucleotide-mediated recombineering followed by <u>B</u>xb1 integrase targeting) system developed by the Sassetti group (1) enables precise gene deletions but has primarily been optimized for *Mycobacterium tuberculosis* and *Mycobacterium smegmatis*. Although ORBIT has been used in Mab, here we describe an optimized ORBIT workflow for efficient gene deletion in Mab. We tested key parameters, including plasmid selection, oligonucleotide design, and recombineering conditions. We evaluated oligonucleotide length and purification to improve recovery of targeted mutants and established conditions that enabled reproducible deletion of multiple target genes. Our findings establish a robust and adaptable ORBIT protocol tailored to the genetic characteristics of Mab. This optimized workflow provides a practical framework for precise genetic manipulation and can facilitate functional studies of genes that contribute to Mab physiology, antimicrobial resistance, and pathogenesis.

**IMPORTANCE:** *Mycobacterium abscessus* causes the majority of intractable infections attributable to members of the *M. abscessus* complex: *M. abscessus, M. massiliense* and *M. bolletii. M. abscessus* infections most commonly present as chronic lung disease or as skin and soft tissue infections, and they are among the most difficult bacterial infections to eradicate, often persisting despite aggressive, prolonged treatment underscoring the need for reliable genetic tools to dissect the biology underlying pathogenesis. The ORBIT system enables precise, markerless gene deletion in mycobacteria, but its application to *M. abscessus* has been limited. Here we present an optimized ORBIT workflow tailored specifically to *M. abscessus* that will facilitate studies on the genes contributing to physiology and pathogenesis of this important pathogen.

## OBSERVATION

*M. abscessus* (Mab) is a rapidly growing nontuberculous mycobacterium associated primarily with chronic pulmonary infections, particularly in individuals with cystic fibrosis or other underlying lung diseases (2, 3). Mab is difficult to treat, with prolonged, multi-drug treatment regimens often failing because of its intrinsic resistance (4-10). Understanding the genetic basis of intrinsic resistance and other clinically relevant phenotypes require efficient methods for targeted genome manipulation.

Several approaches have been used for genetic manipulation of mycobacteria, including homologous recombination, transposon mutagenesis, and CRISPR-Cas9 (11-15). Although unmarked gene deletions in Mab have most commonly been generated using recombineering (16), we evaluated the newer <u>o</u>ligonucleotide-mediated recombineering followed by <u>B</u>xb1 integrase targeting (ORBIT) method developed for and tested in *M. tuberculosis* and *M. smegmatis* (1). An overview of the published ORBIT protocol used for gene deletion developed by the Sassetti group is summarized in Fig. 1. Briefly, an ORBIT oligonucleotide containing an *attP* site is incorporated into the target locus and subsequently recognized by RecT annealase and Bxb1 integrase, which facilitates insertion of a nonreplicating payload plasmid containing the corresponding *attB* site. The payload plasmid carries a selectable marker that can be used to generate chromosomally tagged genes, genes deletions or promoter replacements (1).The application of ORBIT in Mab remains limited (17-19), and a practical framework addressing species-specific parameters affecting its efficiency is lacking. Here, we address the importance of selectable marker choice, oligonucleotide design and purification, mutant confirmation, and target-locus architecture that improved recovery and identification of deletion mutants.

**Fig 1.**
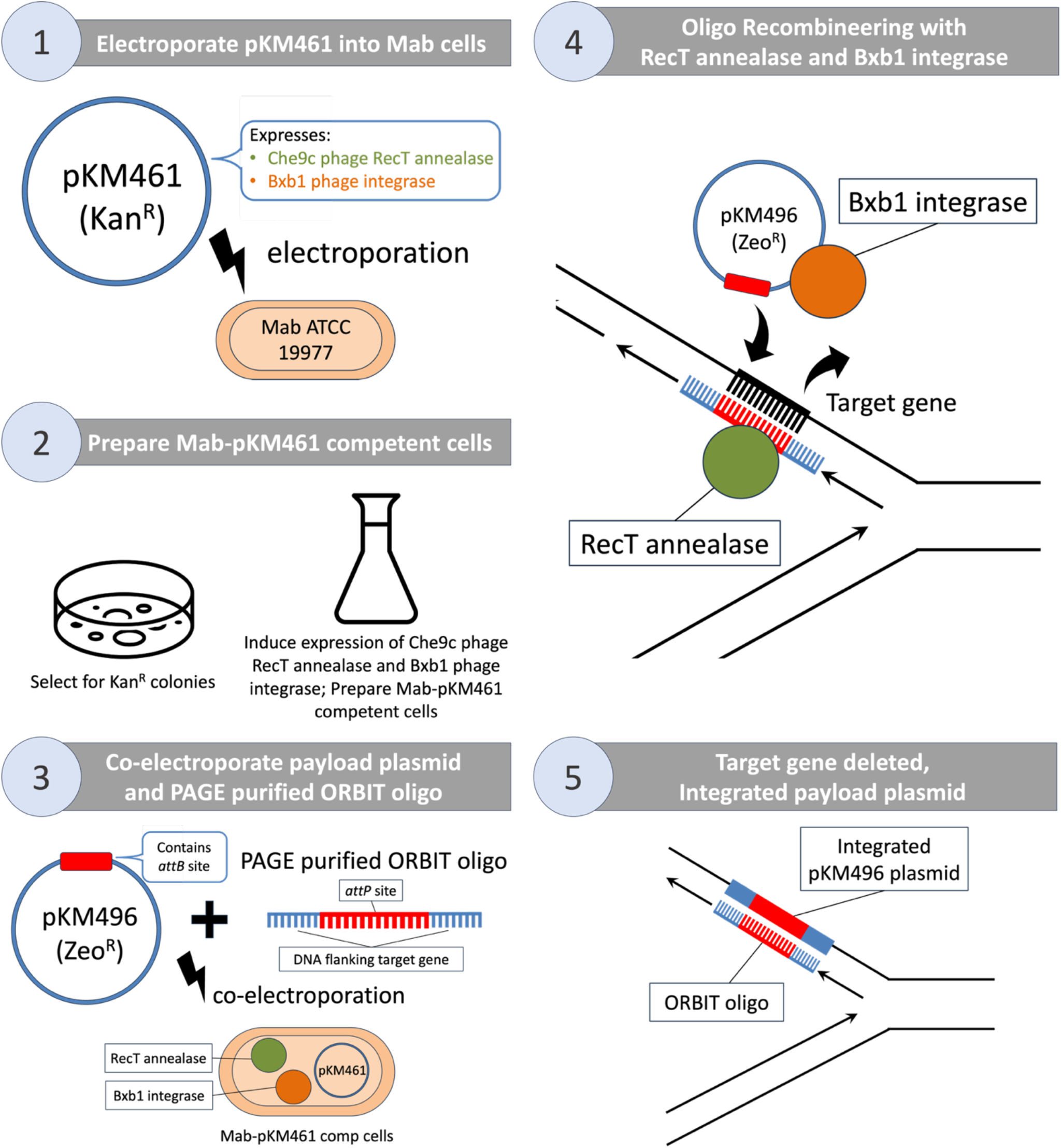
Overview of ORBIT-mediated gene deletion in *Mycobacterium abscessus* based on Murphy et al. (1). 1) The pKM461 plasmid, encoding Che9c phage RecT annealase and Bxb1 phage integrase, is electroporated into Mab ATCC 19977 cells. 2) Transformants harboring the pKM461 plasmid are selected (due to the Kan^R^ selection marker) and competent cells are prepared. 3) The pKM496 payload plasmid, which contains an *attB* site is co-electroporated with a PAGE-purified ORBIT oligonucleotide, which contains an *attP* site flanked by sequences homologous to regions surrounding the target gene. 4) RecT-mediated recombineering replaces the target gene with the *attP* site, enabling Bxb1-mediated integration of the payload plasmid. 5) The resulting mutant contains the integrated payload plasmid in place of the target locus, generating a gene knockout. pKM461 was a gift from Kenan Murphy (Addgene plasmid # 108320 ; http://n2t.net/addgene:108320 ; RRID:Addgene_108320). pKM496 (deletions) Zeo was a gift from Kenan Murphy (Addgene plasmid # 109301 ; http://n2t.net/addgene:109301 ; RRID:Addgene_109301).

### A species-compatible selectable marker is required for ORBIT

We initially attempted to delete the GCN5-related N-acetyltransferase (GNAT) gene *mab_4324c* from the Mab ATCC 19977 genome using the hygromycin-resistant payload plasmid pKM464-Hyg (pKM464 (deletions) was a gift from Kenan Murphy (Addgene plasmid # 108322 ; http://n2t.net/addgene:108322 ; RRID:Addgene_108322). However, we were unable to recover confirmed deletion mutants. To assess hygromycin selection, we plated colonies recovered following co-electroporation of pKM461-containing cells with the ORBIT oligonucleotide and pKM464-Hyg on either hygromycin- or zeocin-containing media. Colonies exhibited confluent lawn growth on hygromycin-containing solid media and substantial growth in liquid media, consistent with intrinsic hygromycin resistance in Mab (20). In contrast, no growth was observed on zeocin-containing solid media, with only minimal growth in liquid media, indicating effective zeocin selection in Mab. We, therefore, replaced pKM464-Hyg with the zeocin-resistant payload plasmid pKM496-Zeo, enabling the recovery of candidate *mab_4324c* deletion mutants. These observations highlight the importance of empirically validating selectable markers for ORBIT in Mab, as markers effective in other mycobacteria species may not provide adequate selection in this organism.

### Longer, PAGE-purified oligonucleotides increase the recovery of correct deletion mutants

We first tested an ORBIT oligonucleotide for *mab_4324c* deletion with a 148-bp Ultramer™ (Integrated DNA Technologies) with 50-bp homology arms flanking the target gene. Because longer homology arms can improve recombineering efficiency (1), we redesigned it as a 188-bp Ultramer™ with 70-bp flanking homology arms. However, longer oligonucleotides yield a lower proportion of full-length product generated during chemical synthesis. Based on IDT’s reported 99.4% per-nucleotide coupling efficiency, the expected full-length yield decreases from ∼41% for a 148-mer to ∼32% for a 188-mer. Therefore, we used PAGE purification to enrich full-length oligonucleotides and incorporated this modification into the published ORBIT protocol to improve recovery of longer oligonucleotides.

Using the 148-mer with 400 ng of the pKM496-Zeo payload plasmid yielded 13 colonies, of which two were confirmed to contain the intended *mab_4324c* deletion (∼15%). In contrast, the 188-mer PAGE-purified oligonucleotide yielded two colonies with 200 ng of pKM496-Zeo and two colonies when used with 400 ng of pKM496-Zeo. Both colonies recovered under the 200 ng condition and one of two colonies recovered under the 400 ng condition contained the intended deletion, resulting in an overall deletion frequency of 75% (3/4 screened colonies; Fig 2A). Thus, although the longer PAGE-purified oligonucleotide yielded fewer colonies, a substantially greater proportion of recovered colonies carried the intended *mab_4324c* deletion.

**Fig 2.**
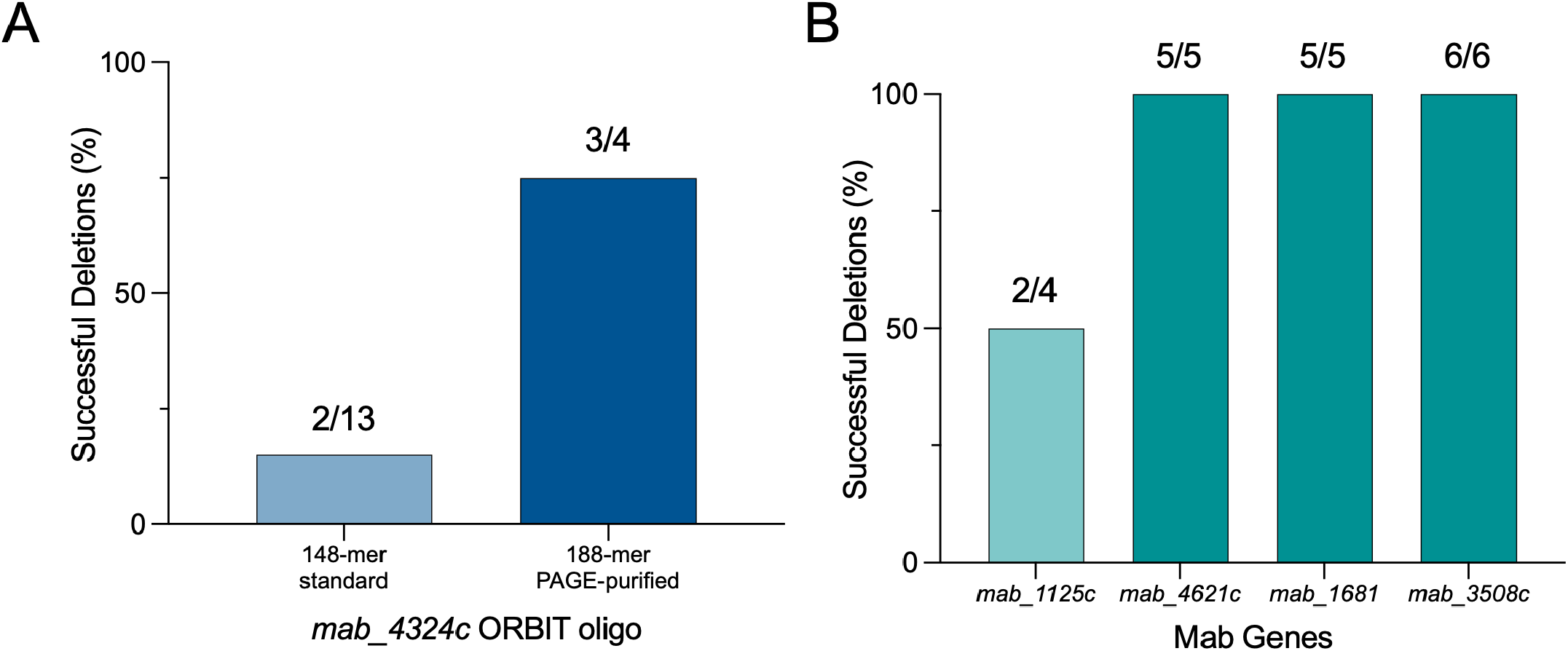
Efficiency of ORBIT-mediated gene deletion in Mab. A) Percentage of successful *mab_4324c* deletion mutants recovered using 4 nmole 148-mer oligonucleotides with standard desalting purification or 4 nmole 188-mer oligonucleotides with PAGE-purification. B) Percentage of successful deletions for additional Mab target genes. *mab_1125c* was targeted using a 4 nmole 148-mer oligonucleotide with standard desalting purification and 200 ng of pKM496-Zeo plasmid. *mab_4621c, mab_1681*, and *mab_3508c* were targeted using 4 nmole 188-mer oligonucleotides with PAGE-purification and 400 ng of pKM496-Zeo plasmid.

### Mab-specific PCR confirmation improves mutant identification

Confirmation primers from the published ORBIT protocol were initially adapted for use with the pKM496-Zeo payload plasmid. However, the original OriE and ZeoR primer combinations produced non-specific amplification in Mab, as these primers had been designed based on *M. tuberculosis* sequences and were not specific for the Mab genome. We therefore redesigned the relevant primers using the Mab ATCC 19977 genome sequence.

To increase confidence in mutant identification, we expanded the PCR confirmation strategy beyond that used in the published ORBIT protocol. Each candidate mutant was screened using multiple independent PCR assays to assess: (i) the presence of the integrated ZeoR cassette, (ii) correct integration of the payload plasmid at either the 5’ or 3’ junctions, (iii) and loss of the target gene. Together, these assays were used to confirm complete, correctly targeted deletion mutants, distinguishing them from colonies containing incomplete or incorrect recombination events, including partial integration of the payload plasmid, ectopic integration, or heterogenous colony populations. Supplemental File 1, Tab 1 lists the PCR confirmation primers and their sequences. Supplemental File 2, Tab 2 provides the expected amplicon sizes for each primer combination used to screen candidate mutant colonies.

### Target-locus architecture must be considered during oligo design

During repeated attempts to delete *mab_4324c*, we identified a 14-bp overlap between the 3’ end of the *mab_4324c* coding sequence and the coding sequence of a neighboring encoded on the opposite DNA strand (Fig. 3A,B). Therefore, complete deletion of *mab_4324c*, including this overlapping 14-bp region, would also remove nucleotides that are part of the neighboring gene and could disrupt its coding sequence. To avoid altering the neighboring gene, we redesigned the ORBIT oligonucleotides so that the 14-bp overlapping region was retained in the chromosome while the remainder of the *mab_4324c* coding sequence was deleted (Fig. 3C,D). Thus, the resulting mutant disrupts *mab_4324c* while preserving the overlapping sequence of the neighboring gene. This finding highlights the importance of evaluating target-locus architecture, including overlapping coding sequences, gene orientation, and neighboring genes, before designing ORBIT oligonucleotides for genome engineering in Mab.

**Fig 3.**
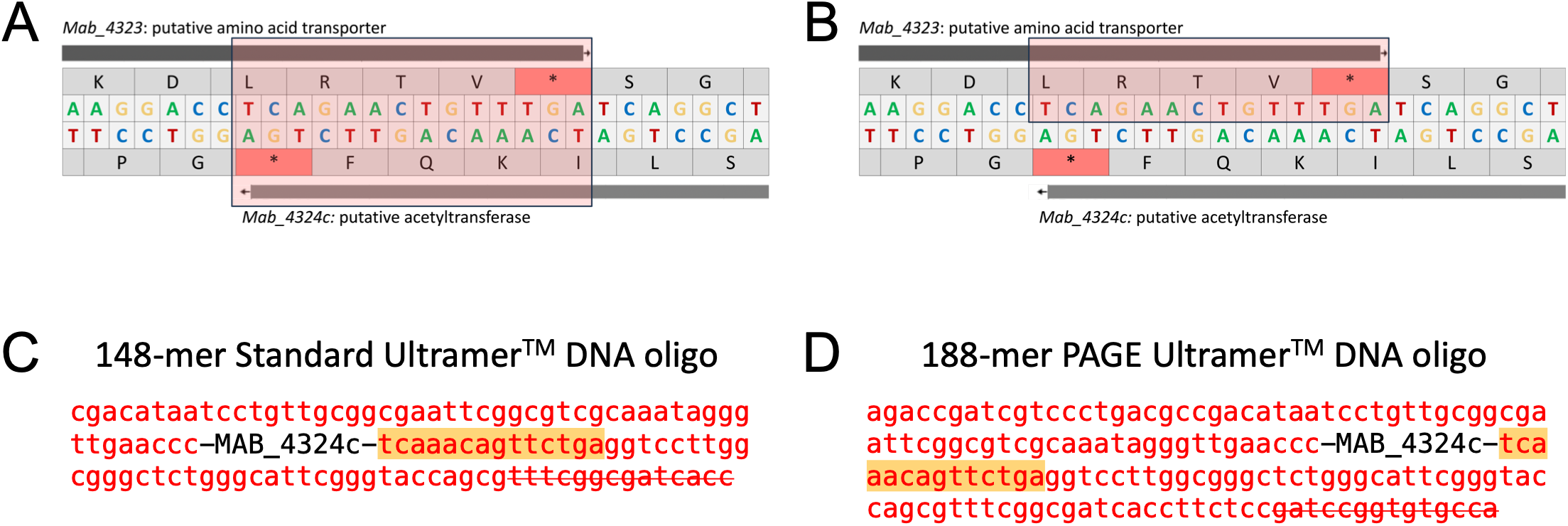
ORBIT oligonucleotide design to preserve an overlapping coding sequence. A) Genomic region showing the 14-bp overlap between the 3’ end of *mab_4324c* and the gene encoded on the opposite strand (pink box). B) The 14-bp overlapping region (pink box) retained as part of the flanking sequence during ORBIT oligonucleotide design. C) Schematic of the 148-mer ORBIT oligonucleotide incorporating the 14-bp overlapping sequence (highlighted in yellow) with the flanking regions (red). D) Schematic of the 188-mer ORBIT oligonucleotide incorporating the 14-bp overlapping sequence (highlighted in yellow) with the flanking regions (red).

### Application of Mab ORBIT to additional Mab loci

To determine whether the optimized ORBIT approach could be applied beyond *mab_4324c*, we targeted three additional GNAT-family genes (*mab_1125c, mab_4621c*, and *mab_1681)* and *whiB7* (*mab_2508c*). The genomic locations and orientations of these target genes in Mab are shown in Fig. 4A, and the lagging strand for ORBIT oligonucleotide design can be identified using the table in Fig. 4B. For *mab_1125c*, we used a 148-mer ORBIT oligonucleotide, and two of four screened colonies were confirmed to contain the intended deletion (Fig. 2B). For *mab_4621c, mab_1681*, and *mab_2508c*, we used longer, PAGE-purified 188-mer ORBIT oligonucleotides; all screened colonies contained the corresponding intended deletion (Fig. 2B). Thus, confirmed deletion mutants were successfully generated for all four additional target genes. Together, these results demonstrate that the optimized ORBIT workflow was successfully applied to multiple loci in Mab. The ORBIT oligonucleotide sequences and PCR confirmation primer sequences are provided in Supplemental File 1, Tabs 3 and 4, respectively.

**Fig 4.**
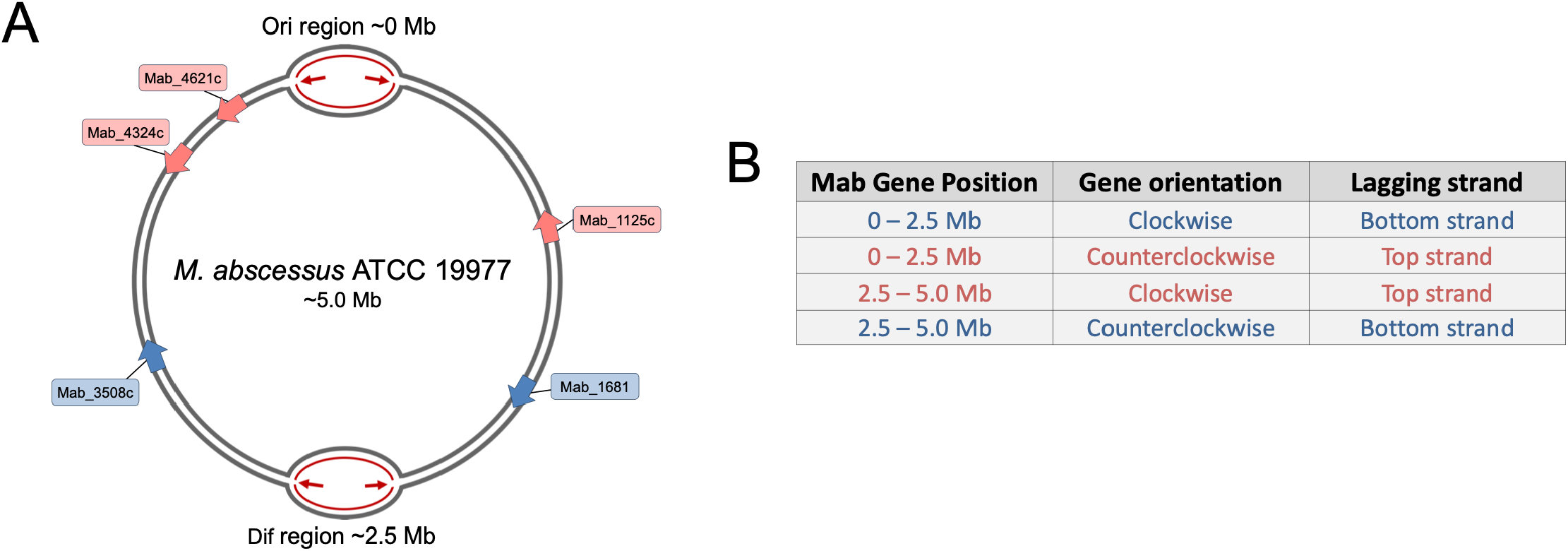
Genomic locations of target genes and lagging-strand identification for ORBIT oligonucleotide design. A) Schematic of the Mab genome showing the locations and orientations of target genes relative to the origin of replication (Ori) and Dif regions. Arrowheads indicate gene orientation; pink and blue denote top- and bottom-lagging strand assignments, respectively, for oligonucleotide design as determined in panel B. B) Table for identifying the lagging strand for ORBIT oligonucleotide design based on genomic position and gene orientation. Blue and pink rows correspond to genes of the same color in Panel A.

## CONCLUSION

The ORBIT protocol by Murphy et al. (1) is a useful approach for targeted gene deletion in Mab but requires species-specific considerations. An Mab-compatible selectable marker was necessary because of intrinsic hygromycin resistance, while longer PAGE-purified oligonucleotides increased the proportion of correctly targeted deletion mutants. Mab-specific confirmation primers and multiple independent PCR assays increased the confidence in mutant identification, and careful evaluation of the target-locus architecture prevented unintended disruption of an overlapping gene. The successful deletion of multiple Mab genes further demonstrates that these modifications can be applied across diverse genomic loci. Together, these findings provide practical guidance for adapting ORBIT-based genome engineering to Mab and establish a framework for future genetic studies of this clinically relevant pathogen.

## ACKNOWLEDGEMENTS

This work was funded by NIH grant R01AI143760 to N.A.W. and T32 GM135141 grant to T.W.S.

## References

1. Murphy KC, Nelson SJ, Nambi S, Papavinasasundaram K, Baer CE, Sassetti CM. ORBIT: a New Paradigm for Genetic Engineering of Mycobacterial Chromosomes. mBio. 2018;9(6).

2. Waugh KM, Wajahat R. Pulmonary Mycobacterium abscessus Infection: A Pathogen in Disguise. Cureus. 2023;15(10):e46897.

3. Abdelaal HFM, Chan ED, Young L, Baldwin SL, Coler RN. Mycobacterium abscessus: It’s Complex. Microorganisms. 2022;10(7).

4. Heid-Picard B, Mougari F, Pouvaret A, Lanternier F, Awad Z, Bille E, et al. Extrapulmonary Mycobacterium abscessus Infections, France, 2012-2020(1). Emerg Infect Dis. 2024;30(11):2294– 302.

5. Jeong SH, Kim SY, Huh HJ, Ki CS, Lee NY, Kang CI, et al. Mycobacteriological characteristics and treatment outcomes in extrapulmonary Mycobacterium abscessus complex infections. Int J Infect Dis. 2017;60:49–56.

6. Hermansen TS, Ravn P, Svensson E, Lillebaek T. Nontuberculous mycobacteria in Denmark, incidence and clinical importance during the last quarter-century. Sci Rep. 2017;7(1):6696.

7. Smith GS, Ghio AJ, Stout JE, Messier KP, Hudgens EE, Murphy MS, et al. Epidemiology of nontuberculous mycobacteria isolations among central North Carolina residents, 2006-2010. J Infect. 2016;72(6):678–86.

8. Brode SK, Marchand-Austin A, Jamieson FB, Marras TK. Pulmonary versus Nonpulmonary Nontuberculous Mycobacteria, Ontario, Canada. Emerg Infect Dis. 2017;23(11):1898–901.

9. Conyers LE, Saunders BM. Treatment for non-tuberculous mycobacteria: challenges and prospects. Front Microbiol. 2024;15:1394220.

10. Yang T, Beach KE, Zhu C, Gan M, Wang W, Zhou H, et al. Genomic Analysis of Global Mycobacterium abscessus Isolates Reveals Ongoing Evolution of Drug Resistance-Associated Genes. J Infect Dis. 2025;231(5):1150–4.

11. Dao TO, Park HE, Lee JH, Kim KM, Trinh MP, Kang HL, et al. Advances and Challenges in Mycobacterial Genetic Engineering: Techniques for Knockout, Knockdown and Overexpression. J Microbiol Biotechnol. 2025;35:e2507051.

12. Chimukuche NM, Williams MJ. Genetic Manipulation of Non-tuberculosis Mycobacteria. Front Microbiol. 2021;12:633510.

13. Alonso MN, Malaga W, Mc Neil M, Jackson M, Romano MI, Guilhot C, et al. Edicient method for targeted gene disruption by homologous recombination in Mycobacterium avium subspecie paratuberculosis. Res Microbiol. 2020;171(5-6):203–10.

14. Pelicic V, Jackson M, Reyrat JM, Jacobs WR, Jr., Gicquel B, Guilhot C. Edicient allelic exchange and transposon mutagenesis in Mycobacterium tuberculosis. Proc Natl Acad Sci U S A. 1997;94(20):10955–60.

15. Li L, Hu Y, Wang D, Li X, Bao S, Deng T, et al. A CRISPR/Cas9-based system using dual-sgRNAs for edicient gene deletion in Mycobacterium abscessus. Front Microbiol. 2025;16:1608274.

16. van Kessel JC, Hatfull GF. Recombineering in Mycobacterium tuberculosis. Nat Methods. 2007;4(2):147–52.

17. Tang D, Wang M, Wang D, Yang D, Cai Y, Luo T, et al. Ultra-high ?eld strength electroporation enables edicient DNA transformation and genome editing in nontuberculous mycobacteria. Microbiol Spectr. 2025;13(10):e0194425.

18. Rodriguez R, Campbell-Kruger N, Gonzalez Camba J, Berude J, Fetterman R, Stanley S. MarR-Dependent Transcriptional Regulation of mmpSL5 Induces Ethionamide Resistance in Mycobacterium abscessus. Antimicrob Agents Chemother. 2023;67(4):e0135022.

19. Jovanovic A, Bright FK, Sadeghi A, Wicki B, Cano Muniz SE, Giannini GC, et al. Large-scale testing of antimicrobial lethality at single-cell resolution predicts mycobacterial infection outcomes. Nat Microbiol. 2026;11(2):566–83.

20. Rominski A, Selchow P, Becker K, Brulle JK, Dal Molin M, Sander P. Elucidation of Mycobacterium abscessus aminoglycoside and capreomycin resistance by targeted deletion of three putative resistance genes. J Antimicrob Chemother. 2017;72(8):2191–200.

